# Long-distance tmFRET using bipyridyl- and phenanthroline-based ligands

**DOI:** 10.1101/2023.10.09.561591

**Authors:** Sharona E. Gordon, Eric G. B. Evans, Shauna C. Otto, Maxx H. Tessmer, Kyle D. Shaffer, Moshe T. Gordon, E. James Petersson, Stefan Stoll, William N. Zagotta

## Abstract

With the great progress on determining protein structures over the last decade comes a renewed appreciation that structures must be combined with dynamics and energetics to understand function. Fluorescence spectroscopy, specifically Förster resonance energy transfer (FRET), provides a great window into dynamics and energetics due to its application at physiological temperatures and ability to measure dynamics on the ångström scale. We have recently advanced transition metal FRET (tmFRET) to study allosteric regulation of maltose binding protein and have reported measurements of maltose-dependent distance changes with an accuracy of ∼1.5 Å. When paired with the noncanonical amino acid Acd as a donor, our previous tmFRET acceptors were useful over a working distance of 10 Å to 20 Å. Here, we use cysteine-reactive bipyridyl and phenanthroline compounds as chelators for Fe^2+^ and Ru^2+^ to produce novel tmFRET acceptors to expand the working distance to as long as 50 Å, while preserving our ability to resolve even small maltose-dependent changes in distance. We compare our measured FRET efficiencies to predictions based on models using rotameric ensembles of the donors and acceptors to demonstrate that steady-state measurements of tmFRET with our new probes have unprecedented ability to measure conformational rearrangements under physiological conditions.

**STATEMENT OF SIGNIFICANCE:** In this work, we expand the working distance of transition metal ion Förster Resonance Energy Transfer (tmFRET) to allow the measurement of donor-acceptor distances from 10 Å to 50 Å. We develop new cysteine-reactive bipyridyl- and phenanthroline-based ligands for Cu^2+^, Fe^2+^, and Ru^2+^ and examine their ability to resolve small ligand-dependent distance changes in maltose binding protein. We extend these studies using pulsed electron paramagnetic resonance spectroscopy to demonstrate the high accuracy of tmFRET for studying protein allostery.

## INTRODUCTION

The rules governing protein function have been intensely studied, with much previous emphasis on structural determination. Our modern understanding of function includes the realization that the static pictures of proteins provided by X-ray crystallography misrepresent the true nature of most proteins as static instead of highly dynamic (1). Indeed, a protein functional state might consist of an ensemble of related structures. The heterogeneity among functionally redundant structures has been explored with techniques like NMR, DEER spectroscopy, and cryoEM, but our ability to map structures onto energy landscapes that explain experimental data is extremely limited. Better approaches are needed to capitalize on the abundance of available structural information and decipher the principles by which proteins function.

Förster resonance energy transfer between a donor fluorophore and an acceptor chromophore was recognized in 1967 by Stryer as a means of measuring distances between donor and acceptor (2) and has since been increasingly used to provide low-resolution structural information on proteins in solution. Typically, pairs of visible-light fluorophores with overlapping emission and absorption spectra are engineered into sites in a protein to report distances between the sites and any changes in distance due to conformational rearrangement of the labeled protein. The efficiency of energy transfer is inversely proportional to the sixth power of the donor–acceptor distance, and FRET can be measured using steady-state fluorescence intensity as either quenching of the donor by the acceptor or sensitization of the acceptor by the donor.

We recently developed a novel system for FRET to improve its ability to resolve state-dependent distance changes in proteins (3–5). We used genetic code expansion to introduce the noncanonical amino acid acridonylalanine (Acd; Figure 1A), which is only slightly bigger than a tryptophan and has no additional linker to the protein backbone (6). As an acceptor, we use colored transition metals, such as Ni^2+^, Co^2+^, and Cu^2+^. Transition metal FRET (tmFRET) confers several advantages compared to standard FRET.

1. Like standard FRET between donor and acceptor fluorophores, tmFRET is steeply distance dependent.
2. tmFRET has little orientation dependence because the metal ion acceptor has multiple transition dipoles (7).
3. The method uses metal binding sites that are small and minimally perturbing, making the position of the metal a more faithful representation of the position of the protein backbone than acceptors with flexible linkers.
4. Metal binding may be rapid and reversible, so the fraction of fluorescence quenched at a saturating concentration of metal gives the absolute FRET efficiency and therefore the distance.
5. In this approach, there is no need to account for spectral contamination of the donor signal by the acceptor, as the acceptor chromophore does not emit light.
6. Metals with different R_0_ values (i.e., different distances producing 50% FRET efficiency) can be chosen to tune the measurement to the distance of interest and distinguish between models more rigorously.
7. Labeling with metal as an acceptor employs chemistry orthogonal to labeling with donor, allowing controlled, site-specific labeling of both sites.
8. Because transition metals absorb light poorly, the donor and acceptor must be very close in order to observe FRET, allowing us to measure short distances (10 – 20 Å), with a steeper distance dependence, than standard FRET (8). This shorter range can be an advantage, allowing small conformational changes to be measured with probes with higher distance resolution, but may also be a limitation when donor–acceptor pairs are more than ∼20 Å apart.

**Figure 1.**
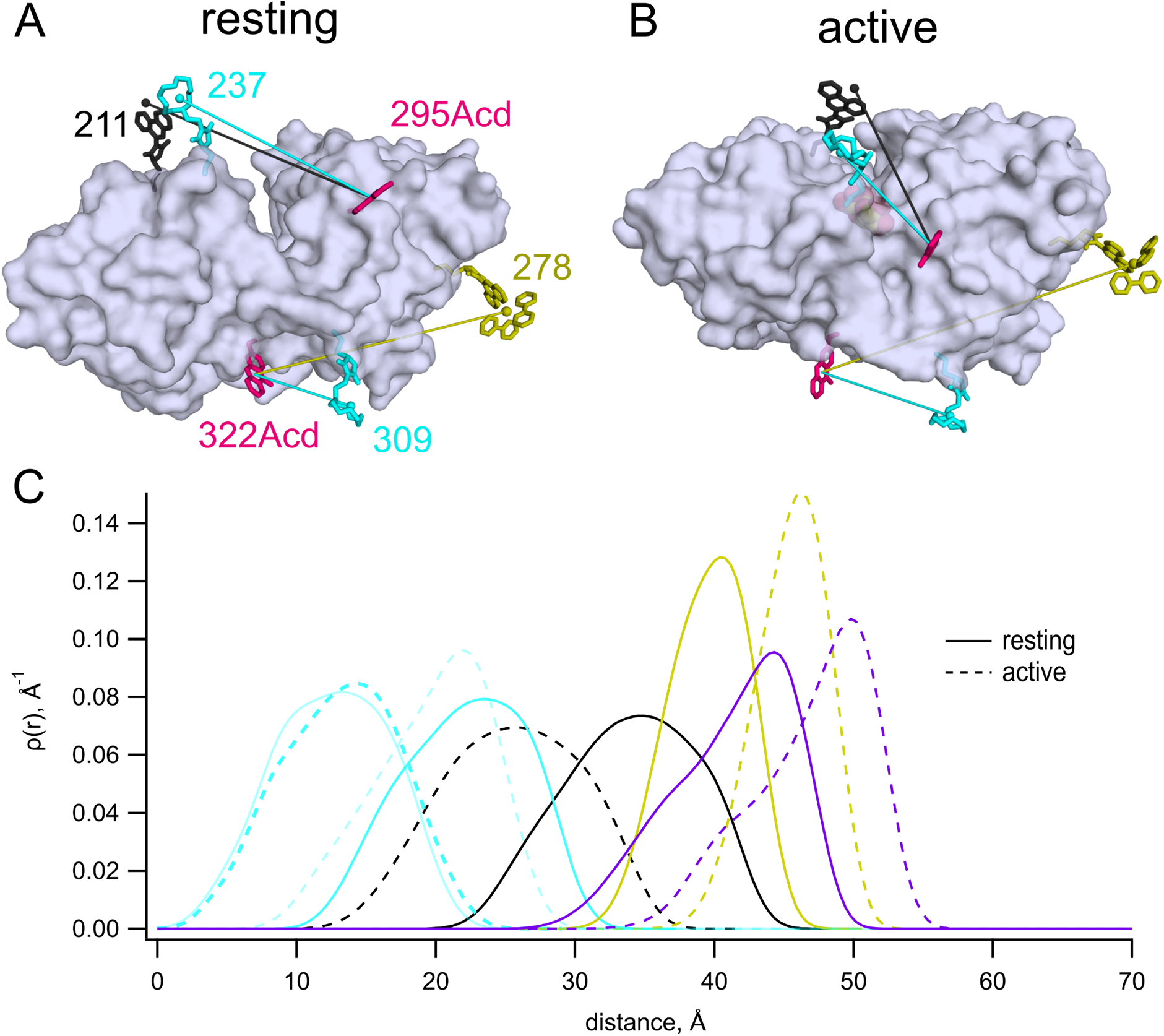
MBP as a model for tmFRET studies. (**A,B**) Cartoon representations of apo (pdb 1omp) (21) (A) and holo (pdb 1anf) (22) (B) MBP labeled *in silico* with 295Acd, 322Acd, 237C-[Cu(TETAC)]^2+^, 309C-[Cu(TETAC)]^2+^, 211C-[Fe(phenM)]^2+^, and 278C-[Ru(bpy)_2_phenM]^2+^. (**C**) Predicted donor–acceptor distance distributions for MBP-295Acd-237C labeled with [Cu(cyclenM]^2+^ (cyan), MBP-322Acd-309C labeled with [Cu(cyclenM]^2+^ (light cyan), MBP-295Acd-211C labeled with [Fe(phenM)]^2+^ (black), and MBP-322Acd- 278C labeled with either [Ru(bpy)_2_phenM]^2+^ (gold) or dabcylM (purple) in the resting (solid) and active (dashed) conformations. Predictions based on chiLife models (14).

Here, we expand the utility of tmFRET to measure ligand-dependent conformational changes across a broader range of distances. We used bipyridyl derivatives, which can bind various transition metal ions to act as tmFRET acceptors that allow us to measure longer distances, in the range of 20-50 Å, due primarily to their greater absorbance of visible light compared to previously used metal-cyclen or metal- histidine complexes. Using maltose binding protein (MBP) as a model system, we measured the steady-state fluorescence of Acd as a donor and various metal-bipyridyls as the acceptor. Our data demonstrate that tmFRET can accurately resolve distances as large as 48 Å. Comparing our measured FRET efficiencies those predicted from structural models of distance distributions demonstrates that, even at the greatest distance between our probes, tmFRET easily revealed small maltose-dependent distance changes. Bipyridyl-metal complexes thus provided a flexible approach to measuring conformational changes across a wide range of distances.

## MATERIALS AND METHODS

### Constructs and Bacterial culture

MBP with a C-terminal twin-strep tag in the pETM11 plasmid and the AcdA9 aminoacyl tRNA synthetase and its cognate tRNA in the pDule2 plasmid (9) were co-transformed into BL-21(DE3) cells using electroporation and grown until single colonies were apparent an plates with appropriate antibiotics as previously described (3). A single colony was grown overnight in 5 mL of Luria Broth and 60 mL of Terrific Broth was inoculated with 60 µL of starter culture and induced when the optical density was 1 with 1 mM IPTG. Acd (0.6 mM in water) was added, and the temperature was then reduced from 37 °C to 18 °C and the cells were then grown overnight. Cell pellets were harvested and stored at −80 °C until use. Note that the MBP-295Acd constructs also included a Y307S mutation to eliminate quenching of Acd by this nearby tyrosine residue (4).

### Protein purification

Cell pellets were resuspended in K^+^-Tris buffer (130 mM KCl, 30 mM Tris, pH 7.4) also containing 2 mM beta mercaptoethanol and cOmplete, EDTA-free protease inhibitor cocktail (Roche Pharmaceuticals, Basil, Switzerland). Cells were lysed by three passes through an Avestin Emulsiflex C3 (Bee International, South Easton, MA). After spinning down cell debris, the supernatant was harvested and passed over Streptactin resin (IBA Life Sciences, Göttingen, Germany) in a disposable column equilibrated with K^+^-Tris buffer. Columns were then washed with 5 column volumes of K^+^-Tris buffer and protein as eluted using 2.5 mM d-desthiobiotin (Sigma-Aldrich, St. Louis, MO) in K^+^-Tris buffer. Bond-Breaker TCEP at 10 mM was added to the appropriate elution fractions and the solution was incubated at room temperature for 10 minutes. This solution was then passed through a PD-10 column (Cytiva Life Science, Marlborough, MA) to remove TCEP and beta mercaptoethanol. Glycerol was added to a final concentration of 10% and the protein was then flash frozen in liquid N_2_ and stored at −80 °C until use.

### Labeling reagents

TETAC, cyclenM, phenM, and [Ru(bpy)_2_phenM]^2+^ were prepared as stocks in DMSO. cyclenM was custom synthesized by AsisChem (Waltham, MA) and [Ru(bpy)_2_phenM]^2+^ was custom synthesized by ACME Biosciences (Palo Alto, CA). TETAC and 1,10-phenM were purchased from Toronto Research Chemicals (Toronto, ON, Canada). All other chemicals and reagents for labeling and fluorescence measurements were purchased from Sigma-Aldrich (St. Louis, MO). 2 M hydroxylamine hydrochloride in water was prepared for use in Fe^2+^ experiments and used for only one day. FeCl_2_ was prepared as a 100 mM stock with 1 M hydroxylamine hydrochloride in water and made fresh for each experiment day.

CuCl2 was prepared as a 100 mM stock in water.

### [Cu(TETAC)]2+ labeling

To label protein with [Cu(TETAC)]^2+^, 100 mM CuCl_2_ in water and 100 mM TETAC in DMSO were mixed at a 1.5:1 ratio and incubated for at least 5 minutes. A 1.5-fold excess of NTA compared to Cu^2+^ was then added to chelate any remaining free Cu^2+^. Water was then added to reach a final concentration of 1.5 mM Cu^2+^, 1 mM TETAC and 2 mM NTA. During the experiment, 2 uL of this solution was added to a total of 200 uL solution in the cuvette, to attain a final concentration of 15 µM Cu^2+^, 10 µM TETAC, and 20 µM NTA.

### [Cu(cyclenM)]2+ labeling

To label protein with [Cu(cyclenM)]^2+^, 100 mM CuCl_2_ in water and 100 mM cyclenM in DMSO were mixed at a 1.5:1 ratio and incubated for at least 5 minutes. A 1.5-fold excess of NTA compared to Cu^2+^ was then added to chelate any remaining free Cu^2+^. Water was then added to reach a final concentration of 12 mM Cu^2+^, 8 mM cyclenM, and 16 mM NTA. During the experiment, 2 uL of this solution was added to a total of 200 uL solution in the cuvette, to attain a final concentration of 120 µM Cu^2+^, 80 µM cyclenM, and 160 µM NTA.

### [Ru(bpy)2phenM]2+ labeling

We prepared a 4 mM [Ru(bpy)_2_phenM]^2+^ stock in DMSO because of the instability of maleimides in aqueous solutions (10–12). During the experiment, we added 1 uL of this solution to a total of 200 µL solution in the cuvette, to attain a final concentration of 20 µM [Ru(bpy)_2_phenM]^2+^.

### [Fe(phenM)3]2+ labeling

For [Fe(phenM)3]^2+^ experiments, a solution of 10.56 mM FeCl_2_ in 105.6 mM hydroxylamine hydrochloride was mixed at a 1:1 ratio with 8 mM phenM in DMSO. After incubating for at least 1 minute, water was then added to bring the final concentration to 1.6 mM FeCl2, 106 mM hydroxylamine hydrochloride, and 1.2 mM phenM. During the experiment, 2 uL of this solution was added to a total of 200 uL solution in the cuvette, to attain a final concentration of 15.6 µM Fe^2+^ and 12 µM phenM. To prevent oxidation of Fe^2+^ to Fe^3+^, 15 mM hydroxylamine hydrochloride was added to protein in a K^+^-Tris buffer at pH 8.3 (Figure S4). After the addition of the hydroxylamine hydrochloride, the pH of the protein-containing solution was 7.4.

### [Fe(phenM)]2+ labeling

For [Fe(phenM)]^2+^ experiments, a 20 mM phenM stock in K^+^-Tris buffer (pH 8.3) was added to MBP protein to achieve a final concentration of 2 mM phenM. After 10 minutes, the solution was passed over a Bio-Rad Micro Bio-Spin 6 column that had been equilibrated with K^+^-Tris buffer (pH 8.3) to remove unreacted phenM label. K^+^-Tris buffer (pH 8.3) was also used to elute the protein. During the experiment, 2 uL of 20 mM Fe^2+^ with 200 mM hydroxylamine hydrochloride was added to a total of 200 µL solution in the cuvette to give a final concentration of 200 µM Fe^2+^ with 2 mM hydroxylamine hydrochloride. The cuvette had an additional 15 mM hydroxylamine hydrochloride, bringing the total concentration to 17 mM. The pH of the solution in the cuvette was approximately 7.4.

### Dabcyl C2 maleimide to correct for incomplete labeling

To ensure complete labeling by the chelator-metal complexes and measure the fraction of cysteines that appear unreactive, we used dabcyl C2 maleimide (dabcylM; Figure 2) as a probe. DabcylM is a non-fluorescent organic quencher with an R_0_ with Acd of 48.9 Å. When used to label MBP-295Acd-237C, MBP-295Acd-211C, and MBP-322-309C, dabcylM quenched at least 92% of the fluorescence, and no change in quenching occurred upon the addition of maltose (Figure S6). These data are consistent with a FRET efficiency near 1 for the 90% of the protein that reacted with dabcylM. When labeling with Cu^2+^- CM, [Ru(bpy)_2_phenM]^2+^, Fe(phenM)^2+^, and [Fe(phenM)_3_]^2+^, we applied dabcylM at the end of the experiment and observed no additional increase in quenching (data not shown). The absence of additional quenching with dabcylM, even for MBP-322Acd-278C, indicates that all reactive cysteines in our protein sample were already labeled with the chelator-metal complexes. The dabcylM experiments also set an upper limit on the unlabeled fraction of our protein at no more than 8%. We used this value to calculate FRET efficiencies from our measured quenching data:

**Figure 2.**
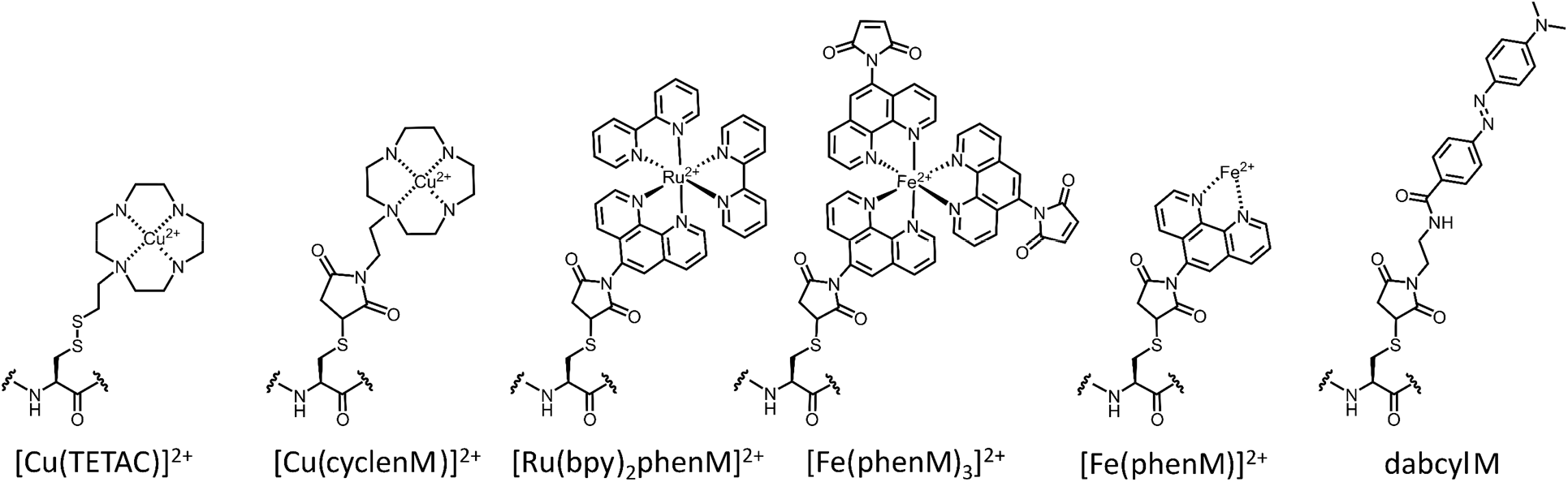
Structure of FRET acceptors used in this study.

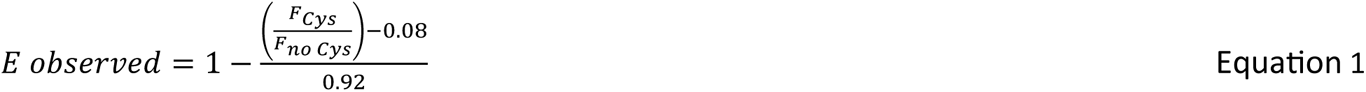

### Spectrophotometry

Absorption measurements were made using a Beckman Coulter DU 800 spectrophotometer (Brea, CA) with a 1 cm path length. For [metal(phen)_3_]^2+^ compounds with Cu^2+^, Co^2+^, and Ni^2+^, we used 1 equivalent of metal to 4 equivalents of phenanthroline, staring with metal stocks made in water and phenanthroline in DMSO. For [Ru(bpy)_3_]^2+^ we prepared a DMSO stock and then diluted into water to achieve the desired concentration. For [Fe(phen)_3_]^2+^, Fe^2+^ which was made in aqueous solution containing a 5-fold excess of hydroxylamine hydrochloride and mixed with phenanthroline stock made in DMSO. The final concentrations used for each metal were: 5 mM CuCl_2_ (20 mM phenanthroline), 20 mM NiCl_2_ (80 mM phenanthroline), 20 µM [Ru(bpy)_3_]^2+^, and 50 µM FeCl_2_ (200 µM phenanthroline). For measuring the spectrum of Fe^2+^ bound to a single phenanthroline, we followed the basic protocol from Kolthoff et al. (13). Briefly, we mixed 2 µL of 5 mM phenanthroline with 98 µL of water, and then added 100 µL of 100 mM Fe^2+^ in 1 M hydroxylamine hydrochloride in water. This gave final concentrations of 50 mM Fe^2+^, 50 µM phenanthroline, and 500 mM hydroxylamine hydrochloride.

### LC-MS methods

Accurate masses of unlabeled and labeled proteins were determined via LC-MS with an Agilent PL1912- 1300 reverse phase HPLC column (Agilent, Santa Clara, CA, USA) connected to an AB Sciex 5600 QTOF mass spectrometer (AB Sciex, Framingham, MA, USA). Mass deconvolution was performed using the Bio Tool Kit microapplication within AB Sciex PeakView 2.2.

### Fluorescence data acquisition and analysis

Data were acquired using a Horiba Fluorolog 3 with an excitation wavelength of 385 nm and an emission wavelength of 425 nm. The excitation and emission slits were always equal and were adjusted between 2.5 and 5 nm, based on the protein concentration, to remain within the linear range of the instrument. Anti-bleaching mode was used to turn off the excitation light between measurements. We used a four- cuvette carousel and interleaved the negative control (protein without the cysteine) with the experimental samples. The workflow for experiments was as follows. Initial measurements in the absence of protein were taken and used for background subtraction. The protein was then added, and measurements were made to establish a baseline for normalization. The metal-chelator complex was then added, and measurements recorded until the fluorescence intensity reached a steady state. Typically, maltose was then added to all cuvettes and, again, the fluorescence intensity was recorded until it reached steady state. For [Cu(TETAC)]^2+^, TCEP was then added to reduce the disulfide bond and remove [Cu(TETAC)]^2+^ from the protein. For all maleimide-reactive chelators, dabcylM was added to ensure that all reactive cysteines were labeled with the metal-chelator complex.

Data were exported to Excel (Microsoft Corp, Redmond, WA) and analyzed as follows. After background subtraction, data were normalized to the initial steady-state fluorescence with protein and no metal.

The repeats of 4-8 experiments using the “F_No Cys_” protein were averaged together and used to normalize the data from the cysteine (F_Cys_) containing protein as F_Cys_/F_no Cys_. Time courses are presented as the mean ± standard error of the mean, with the number of experiments given in the legend of each figure.

### In silico labeling and distance distribution simulations

Computational modeling of Acd and metal-phenM labels, as well as distance distribution predictions, were performed using chiLife (14) with the accessible-volume sampling method (15,16). Acd, and cysteine conjugates of [Ru(bpy)_2_phenM]^2+^, [Fe(phenM)]^2+^, [Fe(phenM)_3_]^2+^, and [Cu(phenM)]^2+^ were added as custom labels in chiLife. Briefly, starting label structures were constructed in PyMOL and energy minimized with the GFN force field in xTB (17). Custom labels were superimposed onto labeling sites of the target pdb structure, and mobile dihedral angles were uniformly sampled. Rotamers with internal clashes (non-bonded atoms < 2 Å apart) were discarded. The energetic contributions of external clashes were modeled using modified Lennard-Jones potential and used to reweight each rotamer as previously described (15). Low weighted rotamers accounting for less than 0.5% of the cumulative ensemble population were discarded from the ensemble. Sampling was terminated once 10,000 samples had been attempted, generating between 400 and 2,500 rotamers, depending on the specific label and protein site. To calculate a simulated distance distribution between two rotamer ensembles, a weighted histogram was made for pairwise distances between the fluorescent centers of each pair of rotamers from the two ensembles. For Acd, the fluorescent center coordinates were defined by the mean position of all atoms in the central acridone ring. For the metal-PhenM labels, the fluorescent center coordinates were on the transition metal ion. Histograms were then convolved with gaussian distributions with a 1 Å standard deviation, and the resulting distributions were normalized such that the integral of the distribution over the distance domain was equal to 1 (P(*r*)).

### FRET efficiency calculation

To calculate the predicted FRET efficiency for a given donor/acceptor pair, the apparent FRET efficiency was calculated using the following equation:

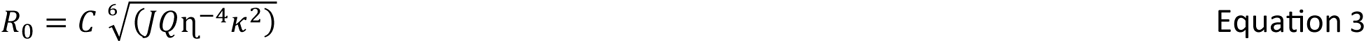

Where *R*_o_values, the distance producing 50% energy transfer, were calculated using the measured donor emission spectrum and quantum yield and the measured absorption spectrum for each type of bound metal using the following equation (18):

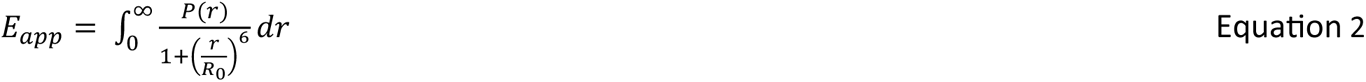

where *C* is a scaling factor, *J* is the normalized spectral overlap of the emission of the donor and absorption of the acceptor, *Q* is the quantum yield of Acd, *ɳ* is the index of refraction (1.33 in our case), and κ^2^ is the orientation factor, assumed to be 2/3. This assumption is justified for at least three reasons. 1) The metal ion acceptor has multiple transition dipole moments. In the case of a mixed polarization donor-acceptor pair, in which one probe is freely rotating and one is immobile, 1/3 < *κ*^2^ < 4/3 (19). Thus, even if our donor were immobile, an assumption of *κ*^2^ = 2/3 gives a maximum error in R_0_ of about ±11%. 2) Our data reveal a distribution of donor-acceptor distances, introducing additional sources of randomized orientation and further reducing the potential error due to the *κ*^2^ = 2/3 assumption. 3) Using *κ*^2^ = 2/3 for our *R_0_* calculations in this and other tmFRET studies gives the expected distances based on known structures.

## RESULTS

### Cu2+-CM is an acceptor for Acd at short distances but not long distances

We have previously used MBP as a model protein for validating new donor–acceptor pairs in tmFRET. MBP is shaped like a clamshell with an equilibrium between its open, or resting, state and its closed, or active, state regulated by sugars such as maltose (Figure 1A,B) (20). Due to its use as an affinity tag in purification, the protein data bank catalogs over 450 MBP structures (search 8/15/2023). MBP lacks native cysteine residues, allowing us to introduce single cysteines to introduce site-specific labels. To probe the conformational rearrangement in MBP, we used amber codon suppression to introduce Acd as a donor and Cu^2+^ bound to the cysteine-reactive chelator cyclen as an acceptor (3,5). Using the molecular modeling software chiLife (14), together with the structures of apo and holo MBP (pdb 1omp and 1anf) (21,22), we modeled the rotameric ensembles of Acd and cysteine-reactive chelators to estimate the donor–acceptor distance distributions in the resting and active states (Figure 1C). We situated one donor–acceptor pair on the lip of the clamshell, with a maltose-induced decrease in *distance* (MBP-295Acd-237C; Figure 1). We used a second donor–acceptor pair on the back side of the clamshell, with a maltose-induced *increase* in distance (MBP-322Acd-309C; Figure 1).

Our previous approach using Cu^2+^ bound to cyclen as a tmFRET acceptor used the cysteine-reactive form of cyclen called 1-(2-pyridin-2-yldisulfanyl)ethyl)-1,4,7,10-tetraazacyclododecane (TETAC; Figure 2), a compound with a 2-mercaptopyridine leaving group that results in a disulfide bond between the protein and the cyclen ring (5). One advantage of TETAC is that a reducing agent can be applied at the end of the experiment to remove the Cu^2+^-bound cyclen and demonstrate that the reaction is fully reversible (Figure S1). When labeling high protein concentrations, however, we found that the reversibility of the disulfide linkage led to thiol exchange with the 2-mercaptopyridine leaving group (Figure S2A). In the present study, we expressed MBP-Acd proteins in bacteria to achieve high concentrations of protein and large fluorescence signals. To avoid the time-dependent replacement of cyclen by 2-mercaptopyridine, we developed a new compound with a maleimide linkage to the cyclen ring (cyclenM; Figure 2). cyclenM reacted with our protein to give irreversible addition of the metal-binding cyclen ring (Figure S2B) (23). An additional advantage of cyclenM is that it can react with the protein in the presence of the reducing agent TCEP, expanding its utility to proteins that, unlike MBP, contain native cysteines and require reducing agents to retain full function.

We measured the fluorescence due to Acd in our protein (either MBP-295Acd-237C or MBP-322Acd- 309C) and then added Cu^2+^-cyclen maleimide ([Cu(cyclenM)]^2+^). We performed parallel experiments with a negative control protein that incorporated Acd at the same sites but had no cysteine. For the cysteine-containing proteins, we observed a decrease in steady-state Acd fluorescence due to quenching by [Cu(cyclenM)]^2+^ reacting with the acceptor site cysteine (Figure 3). For MBP-295Acd-237C, the addition of maltose further increased quenching (Figure 3A,E), as the distance between donor and acceptor decreased with clamshell closure. In contrast, for MBP-322Acd-309C, adding maltose produced an increase in Acd fluorescence intensity (Figure 3B,F), as closing of the clamshell increased the distance between donor and acceptor and gave less FRET. Even though the maleimide-linked cyclen is slightly larger than the disulfide-linked cyclen produced by TETAC, the amplitudes of the tmFRET signals we observed for both MBP-295Acd-237C and MBP-322Acd-309C were similar and the maltose-dependent changes in the tmFRET signal were also similar in [Cu(cyclenM)]^2+^ and [Cu(TETAC)]^2+^ experiments (Figure S1).

**Figure 3.**
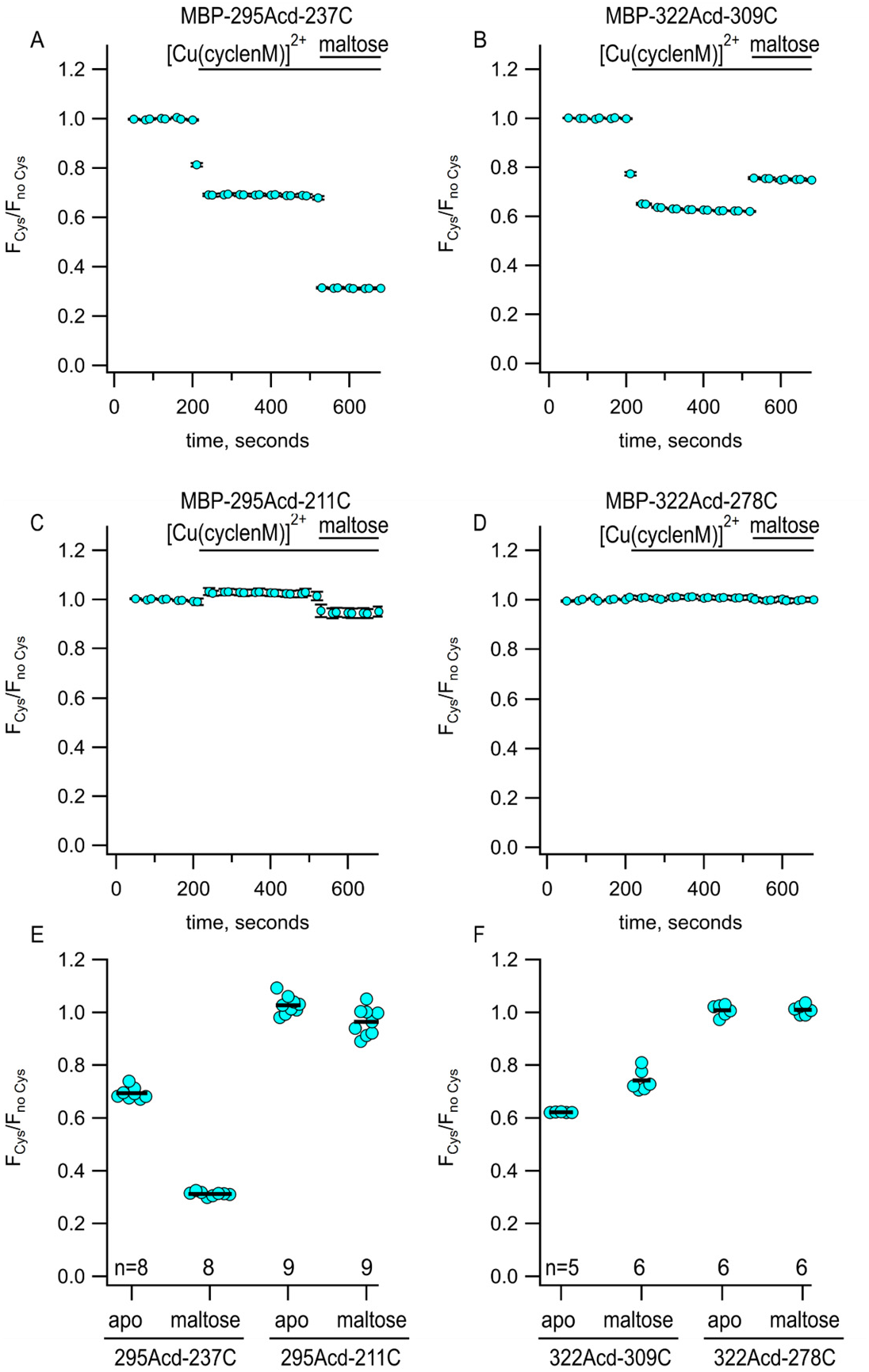
Using Acd and Cu^2+^-CM to measure maltose-dependent change in distance on the lip of MBP clamshell. (**A-D**) Time courses of quenching by Cu^2+^-cyclenM in the absence and presence of maltose, as indicated above each graph. The donor and acceptor site for each experiment are shown above each graph. The number of experimental replicates is listed on the scatter plots in (E) and (F). (**E and F**) Collected data for each of the graphs shown in (A-D) measured at steady state.

The average predicted distances for the two donor–acceptor pairs we examined fall between 13 and 24 Å in both the resting and active states of MBP (Figure 1C, cyan). To determine whether the Acd- [Cu(cyclenM)]^2+^ donor–acceptor pair could be used to probe longer distances, we paired 295Acd with a cysteine at position 211 and 322Acd with a cysteine at position 278. The average predicted distances for these donor–acceptor pairs fall within about 25 to 46 Å (Figure 1C, black and gold). The donor–acceptor distance that produces 50% FRET efficiency (*R*_0_) was calculated to be 15.6 Å for Acd and [Cu(cyclen)]^2+^ (see Materials and Methods). Unsurprisingly, therefore,[Cu(cyclenM)]^2+^ produced little FRET for either MBP-295Acd-211C (Figure 3C,E) or MBP-322Acd-278C (Figure 3D,F), either in the absence or presence of maltose. We concluded that a longer *R*_0_ value would be required to measure these longer distances.

### Bipyridyl-derivative metal complexes provide a flexible platform for tmFRET

We searched for other chelator-metal complexes that might serve as good acceptors in tmFRET experiments. Bipyridyl (bpy)-metal and phenanthroline (Phen)-metal complexes are weakly fluorescent and have been used previously as both FRET donors (24–28) and acceptors (29). As shown in Figure 4A, Ni^2+^, Cu^2+^, Co^2+^, Ru^2+^, and Fe^2+^ all form colored complexes in solution at a 1:3 ratio with bpy or Phen. We measured absorption spectra (Figure 4B) of 1:3 [metal(Phen)_3_]^2+^ complexes for Ni^2+^, Cu^2+^, Co^2+^, Ru^2+^, and Fe^2+^ and 1:1 [metal(Phen)]^2+^ complexes for Cu^2+^ and Fe^2+^ and calculated *R*_0_ values for FRET with Acd ranging from 14 to 44 Å (Figure 4C). We focused on bpy/Phen complexes with Ru^2+^ and Fe^2+^, as these acceptors yielded *R*_0_ values of 24 to 44 Å, allowing us to use tmFRET to measure the longer distances in MBP-295Acd-211C and MBP-322Acd-278C.

**Figure 4.**
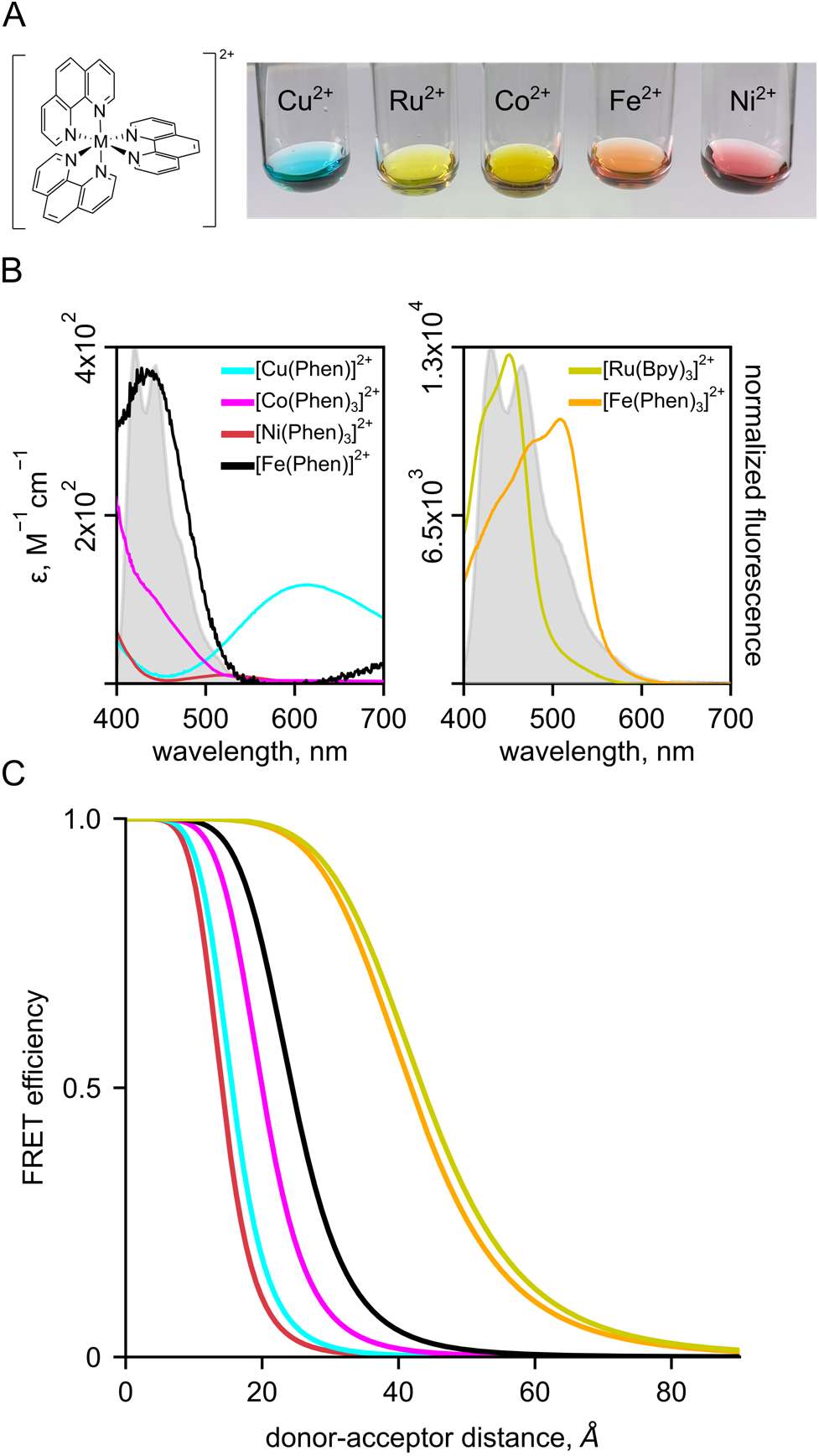
Bipyridyl-metal complexes offer a useful platform for tmFRET. (**A**) Cartoon of Phenanthroline and colored balls representing Ni^2+^, Cu^2+^, Co^2+^, Ru^2+^, Fe^2+^ (3:1 complexes for all), as icons over Photo of Phenantholine bound to different metals. (**B**) Absorption spectra for metal-trisphenanthrolines and the monophenanthroline complex with Fe^2+^. (**C**) FRET efficiency as a function of distance (Forster curves) for phenanthroline metal pairs.

### Using metal-bpy complexes to measure long donor–acceptor distances

We tested if bpy/phen complexes with Ru^2+^ could be used as long distance tmFRET acceptors for MBP- 322Acd-278C. To label MBP with a Ru^2+^-bipyridyl complex, we synthesized the cysteine-reactive [Ru(2,2ʹ- bpy)_2_(1,10-phenanthroline-5-maleimide)]^2+^ ([Ru(bpy)_2_phenM]^2+^; Figure 2), previously used by Lakowicz and colleagues as a rotational probe due to its >1 µs fluorescence lifetime (30). After measuring the initial fluorescence intensity of MBP-322Acd-278C, we added [Ru(bpy)_2_phenM]^2+^ and, after reaching steady-state, added maltose. In the absence of maltose, [Ru(bpy)_2_phenM]^2+^ gave just over 50% quenching and in the presence of a saturating concentration of maltose it gave just under 50% quenching (Figure 5A,B). The calculated *R*_0_ for Acd-[Ru(bpy)_2_phenM]^2+^ of 43.5 Å is thus nicely sandwiched between the resting state and active state distances for MBP-322Acd-278C. Completeness of labeling was evaluated by mass spectrometry (Figure S3) and adding a broad spectrum quencher (dabcyl maleimide; Figure S6) at the end of each experiment (data not shown).

**Figure 5.**
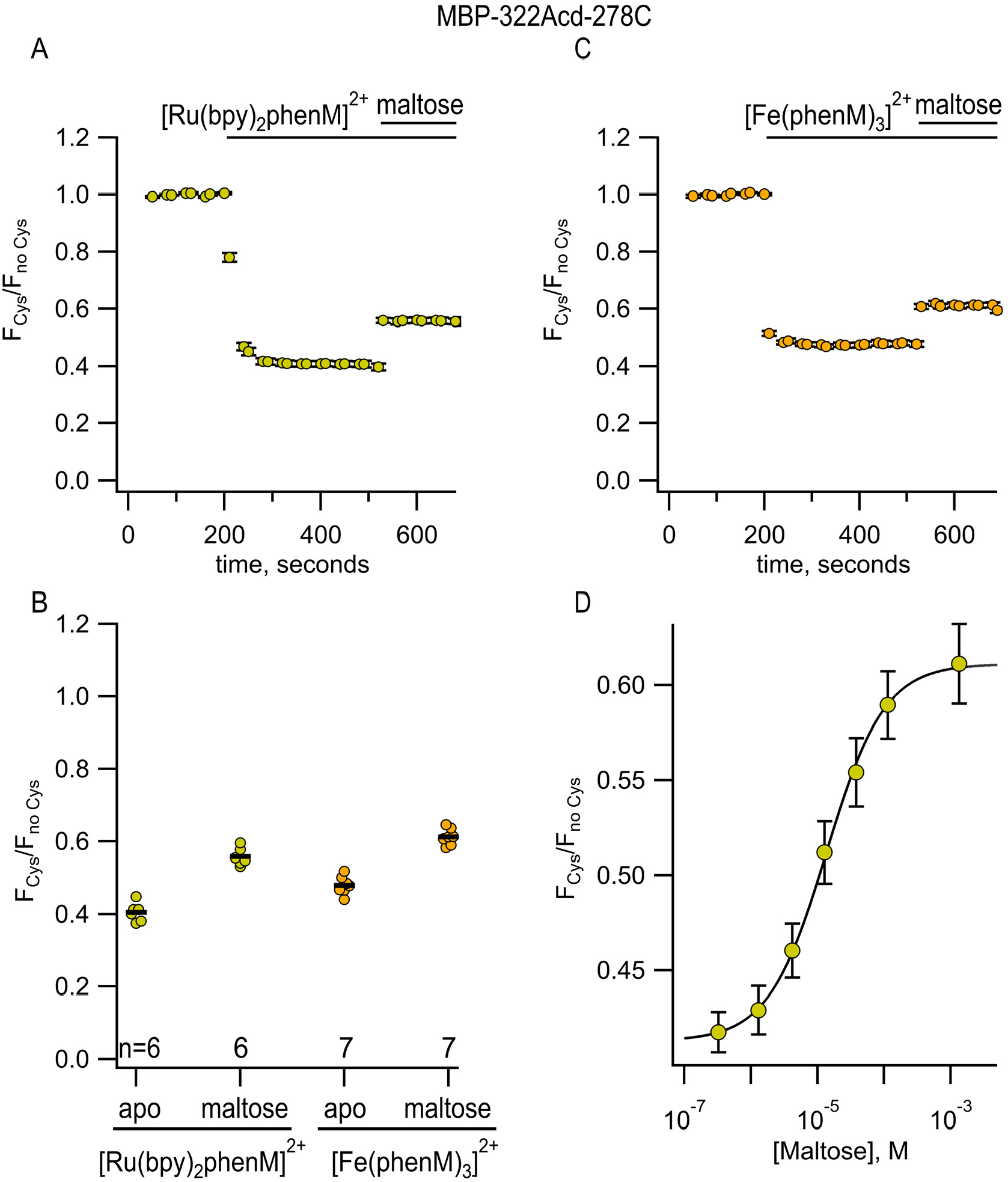
[Ru(bpy)_2_phenM]^2+^ and [Fe(phenM)_3_]^2+^ as tmFRET acceptors for MBP-322Acd-278C, sites on the back side of the MBP clamshell. (**A**) Time course of quenching by [Ru(bpy)_2_phenM]^2+^in the absence and presence of maltose. The number of experimental replicates is listed on the scatter plots in (B). (**B**) Collected data for each of the graphs shown in (A) and (C) measured at steady state. (**C**) Time course of quenching by [Fe(phenM)_3_]^2+^in the absence and presence of maltose. The number of experimental replicates is listed on the scatter plots in (B). (**D**) Dose-response relation for maltose measured using [Ru(bpy)_2_phenM]^2+^. n = 6.

We next used [Ru(bpy)_2_phenM]^2+^ quenching of MBP-322-278C to measure the concentration dependence of maltose binding. The maltose dose–response relation shown in Figure 5D was fit with the Hill equation and gave an apparent *K*_D_ for maltose binding of 13.6 µM, slightly higher than the value reported in the literature (31). These data indicate that our mutations and labeling had only small effects on the apparent affinity of ligand binding, and that tmFRET is a useful way to measure apparent affinities.

We also measured tmFRET in MBP-322Acd-278C with phenM complexes with Fe^2+^. Unlike Cu^2+^ (Figure 6A), Fe^2+^ (Figure 6B) binds to Phen in a highly cooperative manner. The first and second Phen bind to Fe^2+^ with a *K*_D_ of 1.6 x 10^-6^ M and 6.3 x 10^-6^ M, respectively, whereas the third Phen binds with a *K*_D_ of 1 x 10^-10^ M (32). Indeed, the affinity of the 1:3 complex of Fe^2+^ with 3 Phen is so high that even the addition of EDTA did not appear to disrupt it (data not shown). To produce [Fe(Phen maleimide)_3_]^2+^ ([Fe(phenM)_3_]^2+^) (Figure 2), we first mixed Fe^2+^ with 3 equivalents of phenM (Figure S5A). After measuring the initial fluorescence intensity of MBP-322Acd-278C, we then added [Fe(phenM)_3_]^2+^ and, after reaching steady-state, added maltose (Figure 5C). The *R*_0_ of [Fe(phenM)_3_]^2+^ with Acd is 41.8 Å, only slightly less than 43.5 Å for the Acd-[Ru(bpy)_2_phenM]^2+^ pair (Figure 4C). As expected from this almost 2 Å difference, we observed slightly less quenching of MBP-278Acd-278C using [Fe(phenM)_3_]^2+^ compared with [Ru(bpy)_2_phenM]^2+^, in both the absence and presence of maltose (Figure 5B,C).

**Figure 6.**
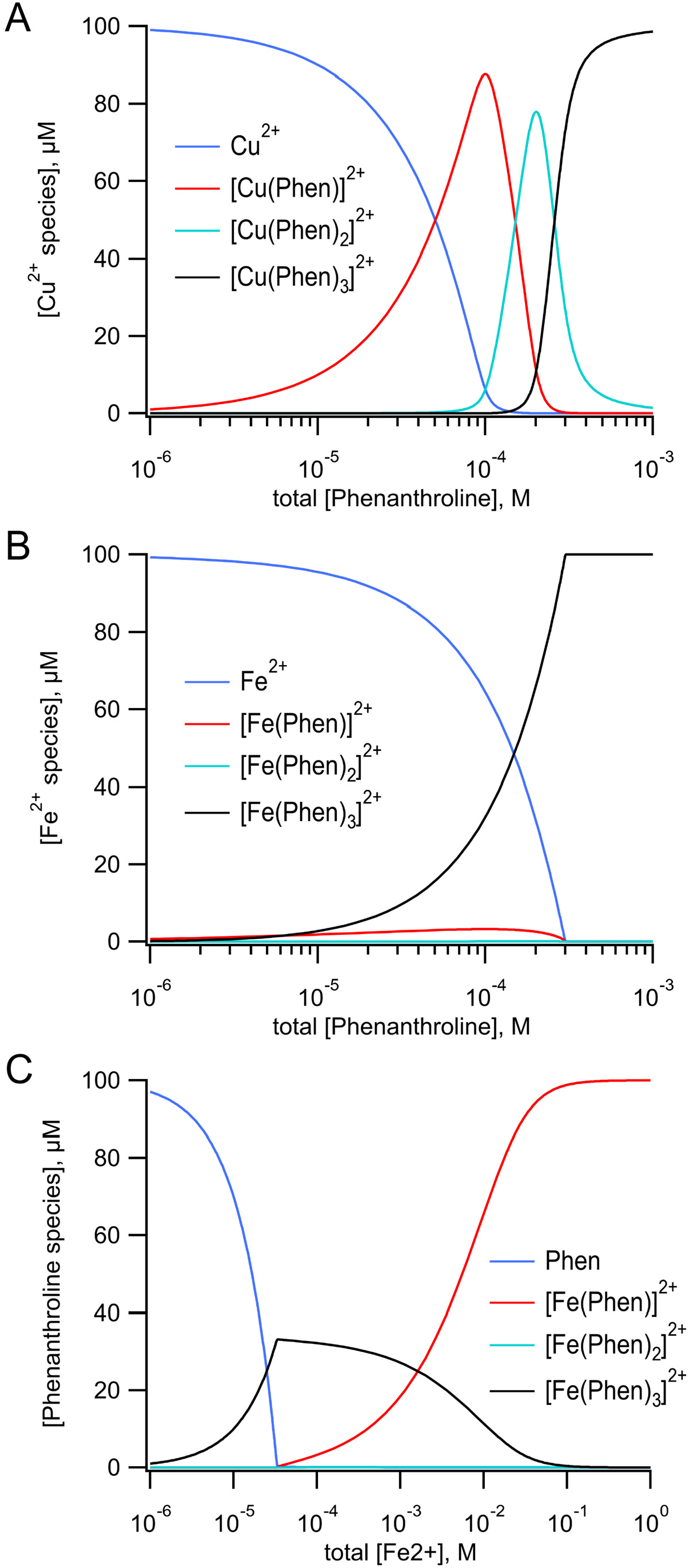
Speciation plots for metal-Phen binding. Thermodynamic model of the binding of three phenanthrolines to a metal ion (top). (**A**) Calculated concentrations of free Cu^2+^, [Cu(Phen)]^2+^, [Cu(phenM)_2_]^2+^, and [Cu(phenM)_3_]^2+^ as a function of the total concentration of Phen with a total concentration of 100 μM Cu^2+^. The values of the association constants K_1_, K_2_, and K_3_ were 1.0 x 10^9^ M^-1^, 5.0 x 10^6^ M^-1^, and 1.0 x 10^5^ M^-1^ respectively (32). (**B**) Calculated concentrations of free Fe^2+^, [Fe(Phen)]^2+^, [Fe(phenM)_2_]^2+^, and[Fe(phenM)_3_]^2+^ as a function of the total concentration of Phen with a total concentration of 100 μM Fe^2+^. The values of K_1_, K_2_, and K_3_ were 6.3 x 10^5^ M^-1^, 1.6 x 10^5^ M^-1^, and 1.0 x 10^10^ M^-1^ respectively (32) .(**C**) Calculated concentrations of free Phen, [Fe(Phen)]^2+^, [Fe(phenM)_2_]^2+^, and [Fe(phenM)_3_]^2+^ as a function of the total concentration of Fe^2+^ with a total concentration of 100 μM Phen. The values of K_1_, K_2_, and K_3_ are the same as in B. The calculations were done using Mathematica (Wolfram Research, Inc.) to simultaneously solve the five equations and five unknowns for the model.

### Using metal-bpy complexes to measure mid-range donor-acceptor distances

We next asked how [Ru(bpy)_2_phenM]^2+^ and [Fe(phenM)_3_]^2+^ would perform for measuring the ∼25 – 35 Å donor–acceptor distances in MBP-295Acd-211C. Consistent with their long *R*_0_ values relative to the donor–acceptor distances at this site, [Ru(bpy)_2_phenM]^2+^ (Figure 7A, D) and [Fe(phenM)_3_]^2+^ (Figure 7B, D) both produced so much quenching that the maltose-dependent change in quenching was difficult to discern.

**Figure 7.**
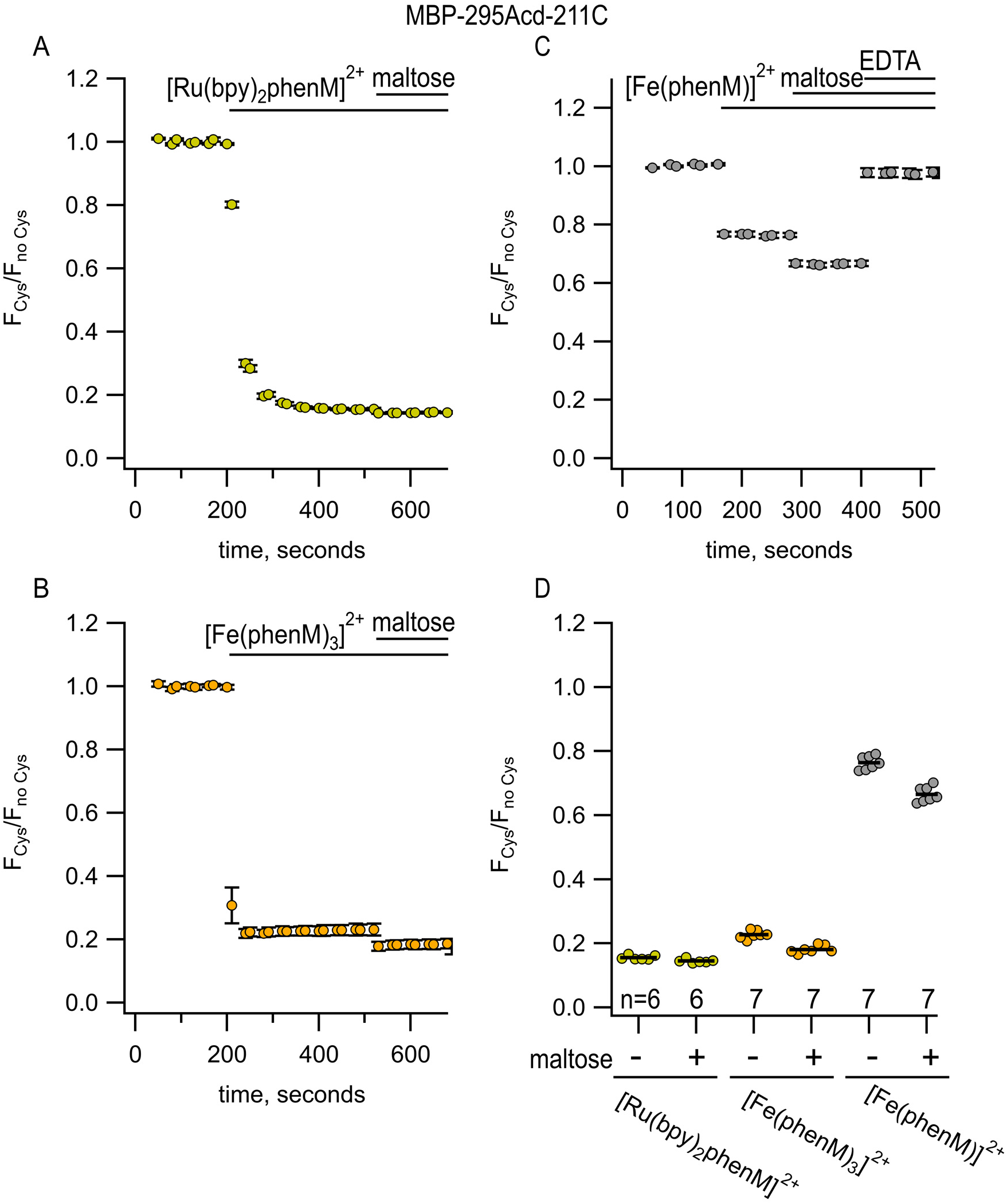
[Ru(bpy)_2_phenM]^2+^, [Fe(phenM)_3_]^2+^, and [Fe(phenM)^2+^ as tmFRET acceptors for MBP-295Acd-211C, sites on the lip of the MBP clamshell. (**A**) Time course of quenching by [Ru(bpy)_2_phenM]^2+^in the absence and presence of maltose. The number of experimental replicates is listed on the scatter plots in (D). (**B**) Time course of quenching by [Fe(phenM)_3_]^2+^in the absence and presence of maltose. The number of experimental replicates is listed on the scatter plots in (D). (**C**) Time course of quenching by [Fe(phenM)]^2+^. (**D**) Collected data for each of the graphs shown in (A-C) measured at steady state.

To better resolve the maltose-dependent change in distance for MBP-295Acd-211C, we used Fe^2+^ bound to just one phenM (Fe(Phen)^2+^; Figure 2), with its smaller *R*_0_ value of 24.4 Å. The high degree of cooperativity in the binding of Phen to Fe^2+^ (Figure 6) required a different labeling strategy than that used in [Fe(phenM)_3_]^2+^ experiments. As depicted in Figure S5B, we first labeled protein with phenM in the absence of metal, column purified the protein to remove unreacted label, and subsequently added a large excess of Fe^2+^ during the experiment to measure fluorescence intensity. The smaller *R*_0_ value of Fe(phenM)^2+^ indeed allowed us to discriminate the maltose-dependent change in distance in MBP- 295Acd-211C (Figure 7C,D).

## DISCUSSION

The Förster equation that relates FRET efficiency to the sixth power of the distance between donor and acceptor assumes a single donor–acceptor distance. This static view of proteins is supported by X-ray crystallography, which generally immobilizes proteins in a single conformational state. In reality, the positions of both the protein backbone and the labels are likely heterogeneous. Instead of a single donor–acceptor distance, then, we should consider a distribution of donor–acceptor distances. We modelled possible donor and acceptor positions using chiLife software as a first approximation of distance heterogeneity (15). ChiLife assumes a static protein backbone and produces a distribution of distances based on predictions of the label rotamer clouds. Donor–acceptor distance distributions predicted by chiLife for several of our sites and labels are shown in Figure 1C. We used the distance distributions predicted by chiLife to calculate predicted FRET efficiencies and plotted these predictions against our observed FRET efficiencies (Figure 8A). The good agreement between our measured efficiencies and those predicted from distance distributions indicates that steady-state FRET measurements accurately reflect donor–acceptor distances if the distance distributions are known or can be approximated for both metal ion acceptors and the small quencher dabcyl. Our predictions of FRET efficiencies might be further improved using structural models that account for motion of the donor and acceptor labels during the lifetime of the fluorophore (33).

**Figure 8.**
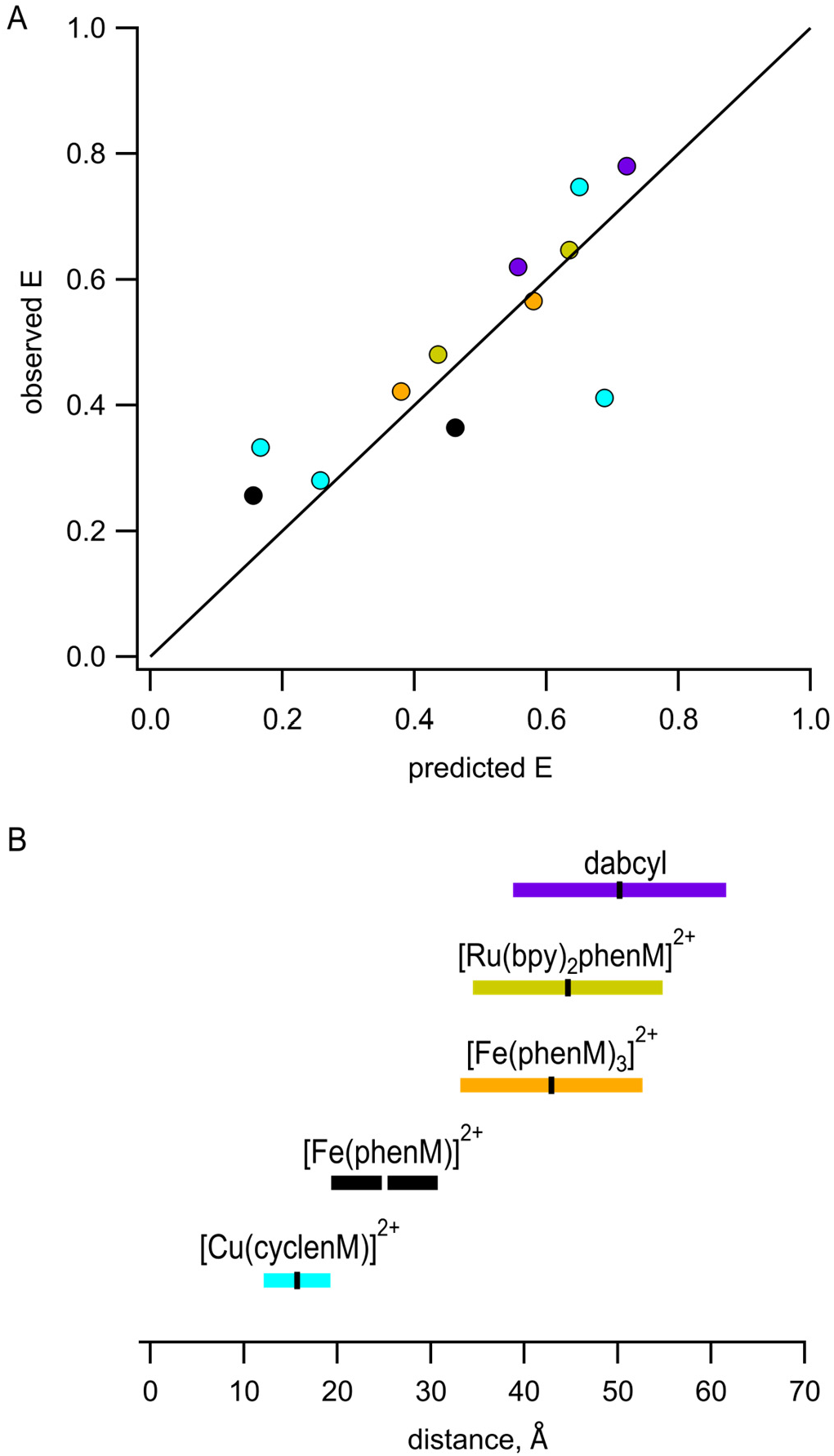
Interpretation of measured FRET efficiences in terms of donor-acceptor distance distributions. (**A**) Comparison of our measured FRET efficiencies with those calculated based on chiLife distance distributions. A line with a slope of 1 is shown for reference. Color schemie is as follows, with R_0_ values in parenthesis: MBP-295Acd-237C and MBP-322Acd-309C labeled with [Cu(cyclenM)]^2+^ - cyan (15.6 Å); MBP-295Acd-211C labeled with [Fe(phenM)]^2+^ - black (24.4 Å); and MBP-322Acd-278C labeled with [Fe(phenM)_3_]^2+^ - orange (41.8 Å), [Ru(bpy)_2_phenM]^2+^ - gold (43.5 Å), or dabcylM – purple (48.9 Å). (**B**) Illustration of the distances for which each of the Acd-acceptor pairs can be measured with tmFRET, defined as the distances that give a FRET efficiency of between 0.8 and 0.2, shown as colored bars, with the corresponding *R_0_* values shown as horizontal ticks.

Building on our previous work using Cu^2+^ bound to cysteine-reactive chelators, which gave a working range for measuring distances of ∼12-19 Å (Figure 8B, cyan; based on distances with predicted FRET efficiencies between 0.8 and 0.2), we focused on expanding the dynamic range of tmFRET to measure distances and distance changes in MBP. Using Fe^2+^ chelated by a single Phen increased the working distance to between 19 Å and 31 Å (Figure 8B, black). Using Fe^2+^ chelated by 3 phenM molecules or Ru^2+^ chelated by one Phen and 2 bpy molecules gave working ranges of 33-53 Å and 35-55 Å, respectively (Figure 8B, orange and gold). Because the *R*_0_, and therefore the working range of FRET, is determined by properties of both the donor and the acceptor, these working ranges could be further expanded using donor fluorophores with different photophysical properties.

In exploring new tmFRET acceptors, we encountered two steps that required significant optimization. Few appropriate (i.e., compact structure with short linkers) cysteine-reactive chelators are commercially available. We designed cyclenM *de novo* and found its irreversible maleimide linkage to cysteines brought a significant advantage compared to TETAC. Although both [Fe(phenM)_3_]^2+^ and [Ru(bpy)_2_phenM]^2+^ use maleimide chemistry to label proteins, they differed in other ways. 1) The [Fe(phenM)_3_]^2+^ complex can be rapidly formed right before the FRET experiment by simply mixing a solution of FeCl_2_ with a solution of phenanthroline maleimide at a ratio of 2.5:1. Because of the highly cooperative nature of the binding of Fe^2+^ to Phen, all of the Fe^2+^ is predicted to form a 3:1 complex without any metal-free phenM to react with the protein (Figure 6B-C; (32)). 2) The *R*_0_ for [Fe(phenM)_3_]^2+^ with Acd is about 2 Å shorter than for [Ru(bpy)_2_phenM]^2+^ (41.8 Å vs. 43.5 Å). However, with more red- shifted fluorophores the differences are more dramatic. For example, with fluorescein, the *R*_0_ values for [Fe(phenM)_3_]^2+^ and [Ru(bpy)_2_phenM]^2+^ are 45.5 Å vs. 32.9 Å and with tetramethylrhodamine (TMR), the *R*_0_ values are 31.4 Å vs. 24.4 Å respectively. This allows for fine tuning FRET pairs to the distances desired. 3) [Fe(phenM)_3_]^2+^ has three maleimide groups and therefore is expected to react more rapidly (or at lower concentrations) and is somewhat bulkier than [Ru(bpy)_2_phenM]^2+^. As shown in a companion paper, however, the extra bulk of these added maleimide groups does not appear to broaden donor– acceptor distance distributions (34). 4) The Fe^2+^ is rapidly oxidized to Fe^3+^ at neutral pH. Thus, these experiments all must be done in the presents of >10 mM hydroxylamine hydrochloride and the measurements made shortly after adding [Fe(phenM)_3_]^2^.

The use of Cu^2+^ also involves significant challenges. Because Cu^2+^ is oxidizing, free Cu^2+^ applied to cysteine-containing proteins induces disulfide bond formation. Ideally, the target cysteine at the acceptor site would be functionalized with cyclenM and then Cu^2+^ would be added. Unfortunately, Cu^2+^ binds to cyclen extremely slowly. Pre-binding of Cu^2+^ to cyclenM at 50 mM each was complete in ∼1 minute. When we then applied Cu^2+^ pre-bound to cyclenM, we did so in the presence of an excess of nitrilotriacetic acid to prevent disulfide bond formation catalyzed by free Cu^2+^ in the solution.

FRET is a better tool for interrogating protein dynamics than for determine protein structure. Compared to X-ray crystallography, cryoEM, and NMR spectroscopy, structural information provided by FRET is sparse and low resolution. The agreement between our measured FRET efficiencies and predicted distance distributions (Figure 8A) is encouraging. However, there are several limitations to FRET that reduce our present ability to extract structural or energetic information from tmFRET data (35). 1) The position of the donor fluorophore and acceptor metal ion may not faithfully report the position of the protein backbone, a problem we have partially mitigated with the use of small donors and acceptors with short linkers to the backbone. 2) The Förster equation assumes a single donor–acceptor distance. 3) Our analysis assumes a single protein conformational state under any given set of conditions. 4) Analysis is complicated by uncertainty about whether there are other pathways for energy transfer (e.g., photoinduced electron transfer), which may be especially problematic at short distances (< 15 Å), especially for aromatic compounds like phenanthroline. 5) Converting measured quenching to FRET efficiency requires measuring or assuming the fraction of protein that is not labeled by acceptor. We determined this unlabeled fraction empirically using the broad-spectrum quencher dabcyl maleimide (Figure S6). 6) The orientation factor *K*^2^may deviate from the 2/3 assumption, though this is less of a concern with isotropic acceptors like transition metals (19). 7) For any given donor–acceptor pair, the range of distances that can be measured is relatively small (Figure 8B). And 8), the distance dependence of FRET is less steep for donor–acceptor pairs with long *R*_0_ values. In the companion paper that follows (34), we will use time-resolved tmFRET to circumvent or mitigate many of these limitations.

## AUTHOR CONTRIBUTIONS

SEG, EGBE, and WNZ designed experiments, performed research, analyzed data, and wrote the paper, SCO and MTG performed research and analyzed data, MHT, EJP, and ST analyzed data, contributed tools, and edited the manuscript, and KDS contributed tools.

## ACKNOWLEDGEMENTS

We thank the Oregon State University GCE4ALL (Center for Genetic Code Expansion for All) for their longstanding collaboration, Dr. Chloe Jones (University of Pennsylvania) for excellent technical support, and Richard W. Aldrich for inspiration throughout our careers. Research reported in this publication was supported by the National Institutes of Health under award numbers R35GM145225 (to S.E.G.), R35GM148137 and R03TR004135 (to W.N.Z.), R01GM125753 (to S.S.), and T32EY007031 (to E.G.B,E). This material is also supported in part by the National Science Foundation under Grant DGE-1747486 (to E.J.P).

---

**Figure S1.**
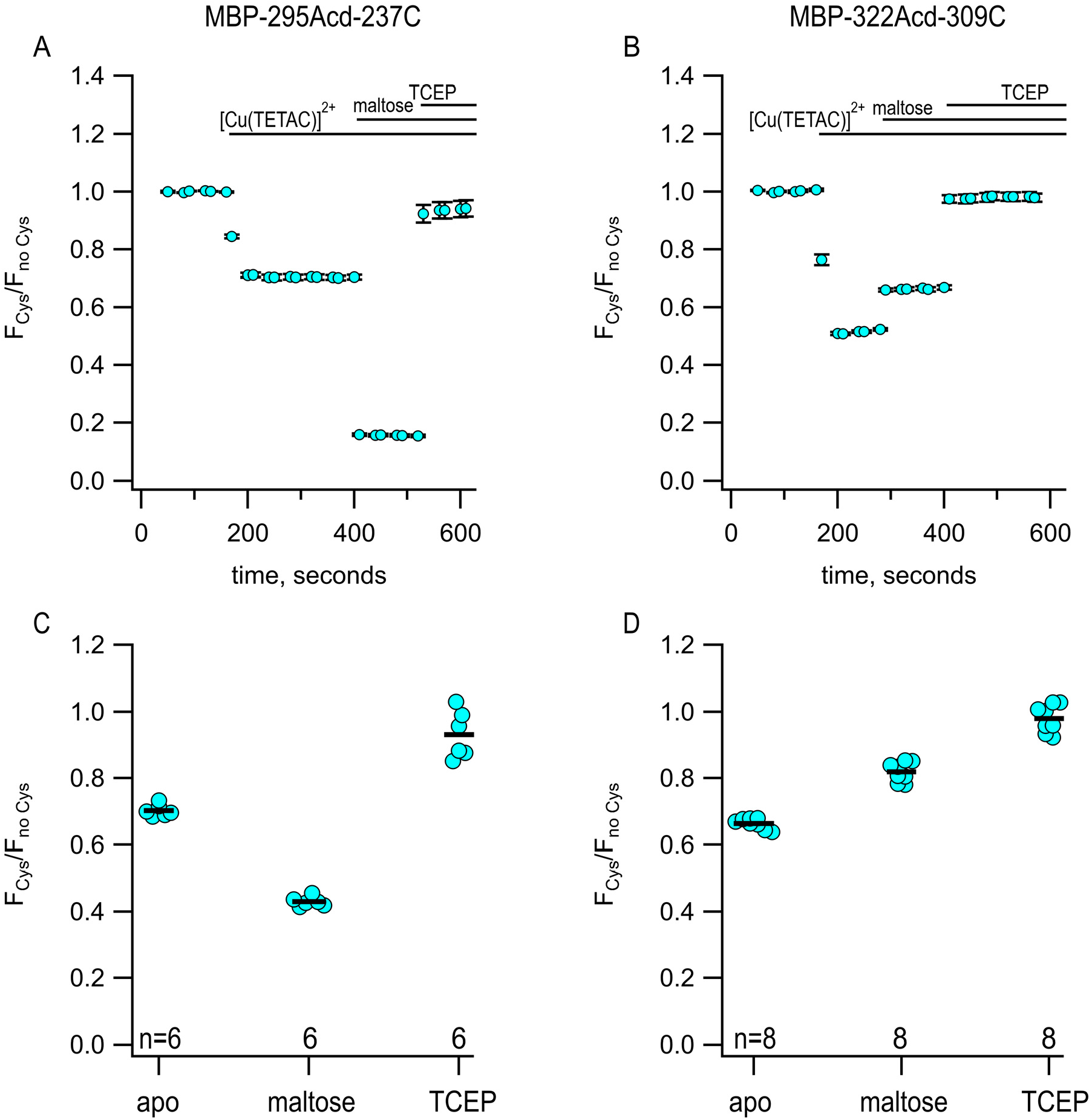
Using Acd and [Cu(TETAC)]^2+^ to measure maltose-dependent change in distance on the lip of MBP clamshell. (**A,B**) Time courses of quenching by Cu^2+^-cyclenM in the absence and presence of maltose, as indicated above each graph. The donor and acceptor site for each experiment are shown above each graph. The number of experimental replicates is listed on the scatter plots in (C) and (D). (**C,D**) Collected data for each of the graphs shown in (A) and (B) measured at steady state.

**Figure S2.**
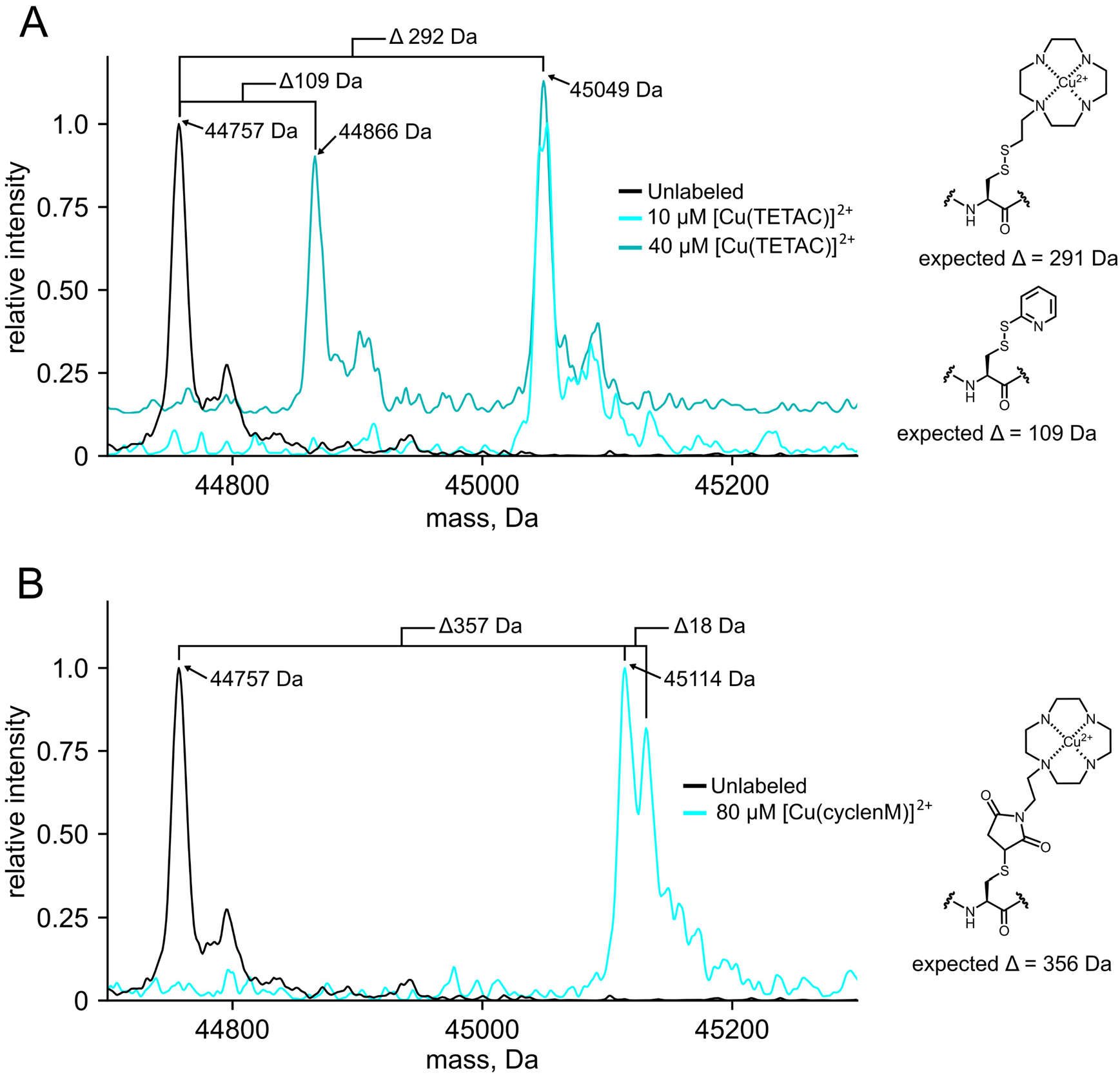
Whole protein mass spectra comparing cyclen chelator labeling strategies. (**A**) Mass spectra of MBP- 295Acd-237C before and after labeling with 10 and 40 μM [Cu(TETAC)]^2+^. The 40 μM [Cu(TETAC)]^2+^ spectrum is vertically offset for clarity. (**B**) Mass spectra of MBP-295Acd-237C before and after labeling with 80 μM cyclen maleimide. The Δ18 peak seen in the labeled sample comes from the hydrolysis of the maleimide ring.

**Figure S3.**
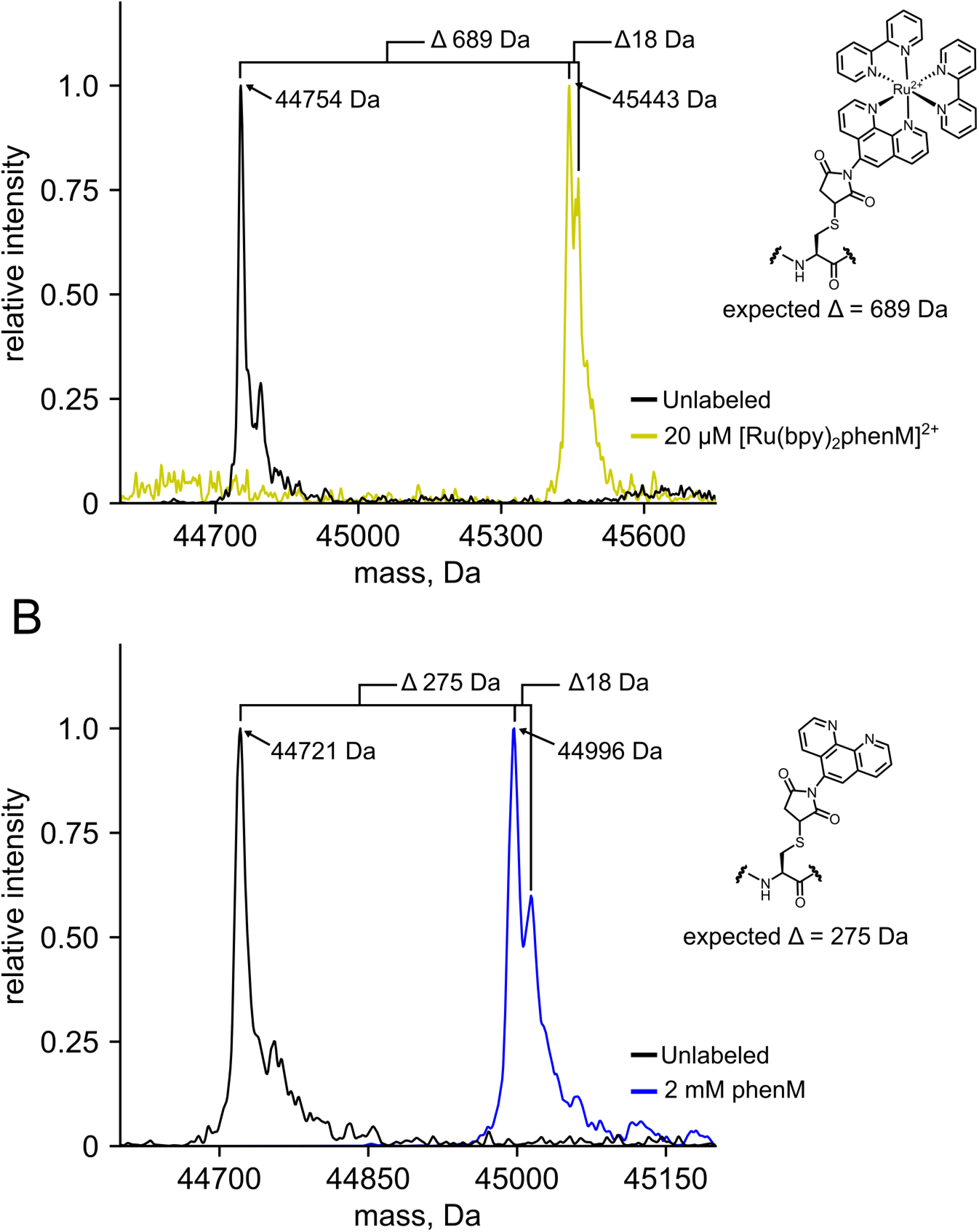
Whole protein mass spectra confirm labeling with metal chelators. (**A**) Mass spectra of MBP-322Acd-278 before and after labeling with [Ru(bpy)_2_phenM]^2+^. (**B**) Mass spectra of MBP-295Acd-211C before and after labeling with phenM. The δ18 peak seen in both labeled samples comes from the hydrolysis of the maleimide ring.

**Figure S4.**
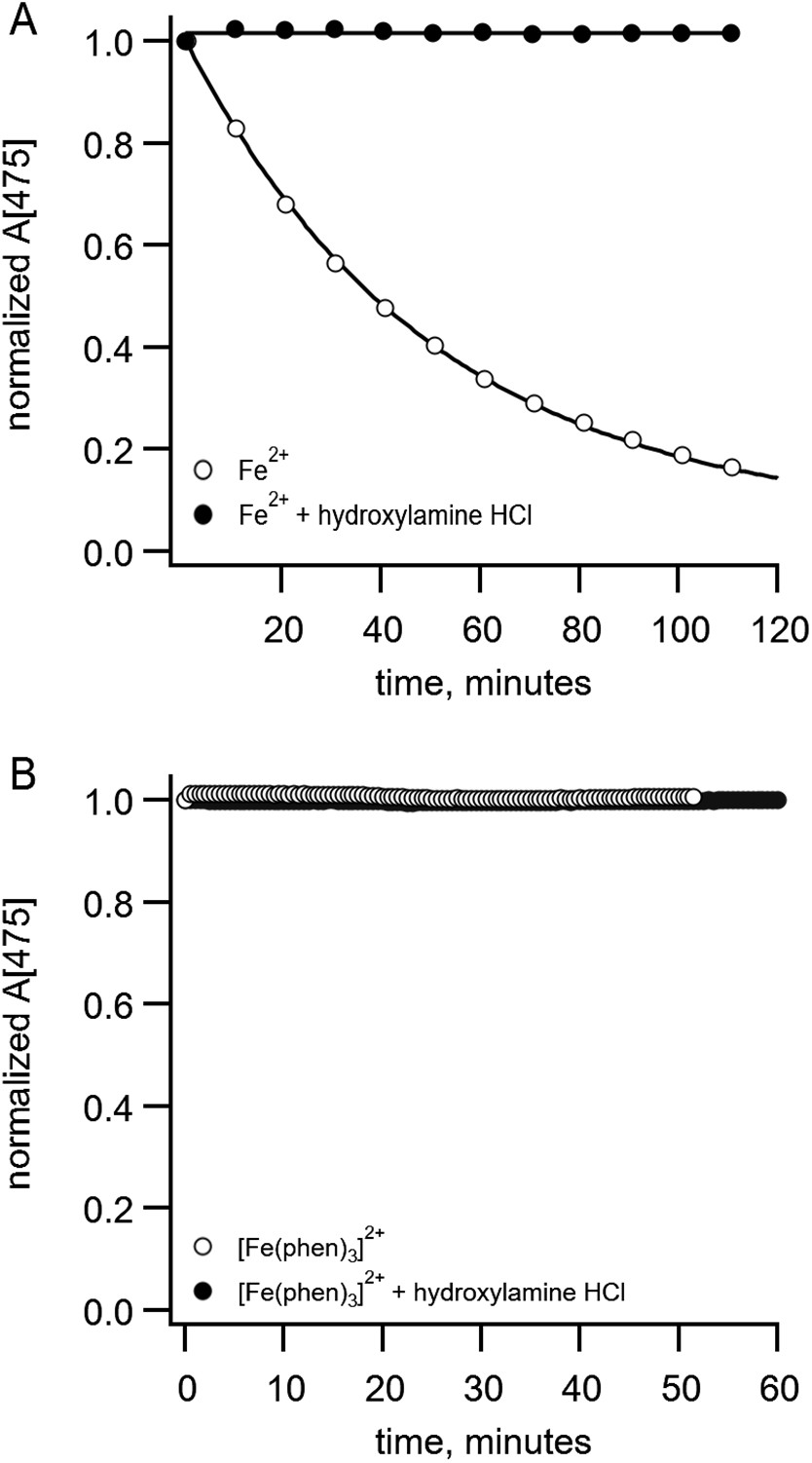
Hydroxylamine hydrochloride inhibits oxidation of Fe^2+^ to Fe^3+^. (**A**) Normalized absorption of 50 µM Fe^2+^ at 475 nm vs time of incubation of in KBT buffer. In the absence (open symbols) or presence (filled symbols) of 15 mM hydroxylamine hydrochloride. The solid curve is a single exponential fit with a time constant of 50 minutes. Phenanthroline (875 µM) was added immediately before each measurement was made. (B) Fe^2+^ bound to phenanthroline did not undergo oxidation even in the absence of hydroxylamine hydrochloride. Normalized absorption at 475 nm vs time of incubation of in KBT buffer for 50 µM Fe^2+^ incubated in KBT in the presence of 875 µM phenanthroline (i.e., [Fe(phen)_3_]^2+^) in the absence (open symbols) or presence (filled symbols) of 15 mM hydroxylamine hydrochloride.

**Figure S5.**
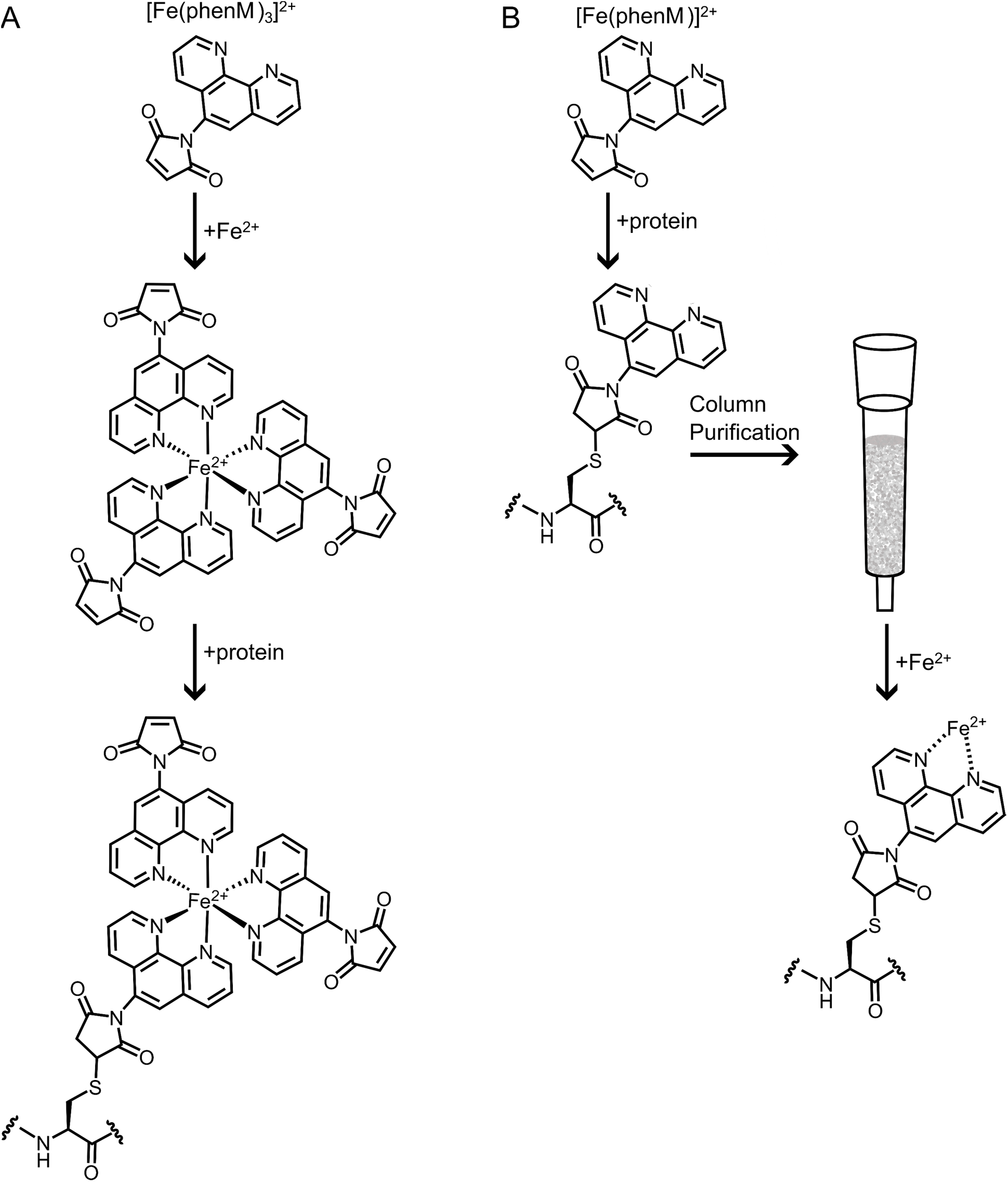
Strategies for labeling with (**A**) 1:3 Fe^2+^:phenanthroline and (**B**) 1:1 Fe^2+^:phenanthroline.

**Figure S6.**
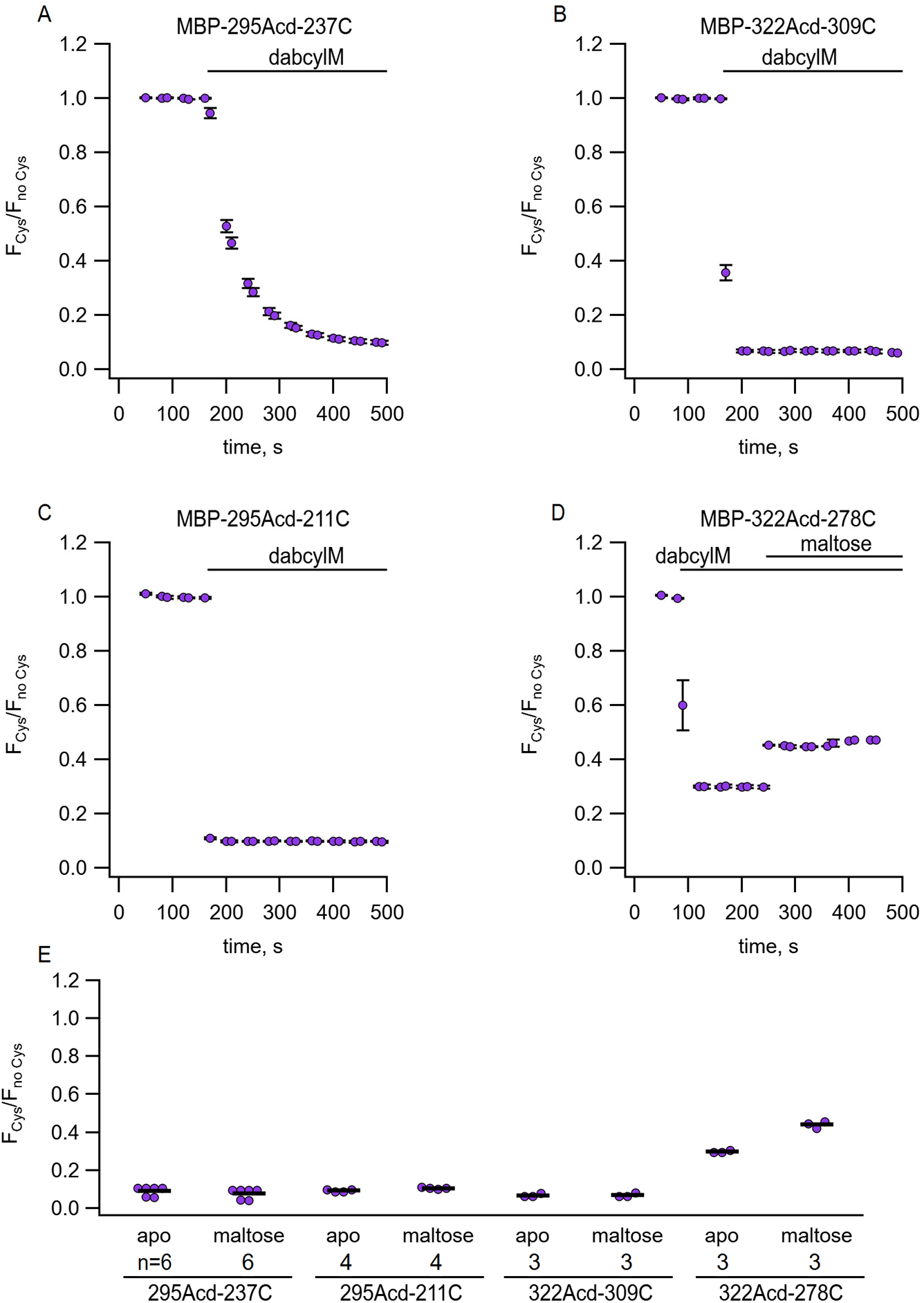
Quenching of Acd in MBP by dabcylM. Time courses of quenching of (**A**) MBP-295Acd-237C; (**B**) MBP- 322Acd-309C; (**C**) MBP-295Acd-211C; and (**D**) MBP-322Acd-278C. The number of replicates for each experiment is shown in (E). (**E**) Collected data for each of the graphs shown in (A-D) measured at steady state.

